# Activation of AKT induces EZH2-mediated β-catenin trimethylation in colorectal cancer

**DOI:** 10.1101/2023.01.31.526429

**Authors:** Ahmed H. Ghobashi, Truc T. Vuong, Jane W. Kimani, Heather M. O’Hagan

**Author notes:** Corresponding author. 1001 East 3^rd^ Street, Bloomington, IN 47405.

## Abstract

Colorectal cancer (CRC) develops in part through the deregulation of different signaling pathways, including activation of the WNT/β-catenin and PI3K/AKT pathways. Enhancer of zeste homolog 2 (EZH2) is a lysine methyltransferase that is involved in regulating stem cell development and differentiation and is overexpressed in CRC. However, depending on the study EZH2 has been found to be both positively and negatively correlated with the survival of CRC patients suggesting that EZH2’s role in CRC may be context specific. In this study, we explored how PI3K/AKT activation alters EZH2’s role in CRC. We found that activation of AKT by PTEN knockdown or by hydrogen peroxide treatment induced EZH2 phosphorylation at serine 21. Phosphorylation of EZH2 resulted in EZH2-mediated methylation of β-catenin and an associated increased interaction between β-catenin, TCF1, and RNA polymerase II. AKT activation increased β-catenin’s enrichment across the genome and EZH2 inhibition reduced this enrichment by reducing the methylation of β-catenin. Furthermore, PTEN knockdown increased the expression of epithelial-mesenchymal transition (EMT)-related genes, and somewhat unexpectedly EZH2 inhibition further increased the expression of these genes. Consistent with these findings, EZH2 inhibition enhanced the migratory phenotype of PTEN knockdown cells. Overall, we demonstrated that EZH2 modulates AKT-induced changes in gene expression through the AKT/EZH2/ β-catenin axis in CRC with active PI3K/AKT signaling. Therefore, it is important to consider the use of EZH2 inhibitors in CRC with caution as these inhibitors will inhibit EZH2-mediated methylation of histone and non-histone targets such as β-catenin, which can have tumor-promoting effects.

## Introduction

Colorectal cancer (CRC) is the third leading cause of cancer-related mortalities in the US (1). CRC develops in part through the deregulation of different signaling pathways, including activation of the WNT/β-catenin and PI3K/AKT pathways (2,3). Activating mutations in the WNT/β-catenin pathway occur in almost 90% of CRCs and are key initiating and promoting events in CRC development (4). Mutations in the PI3K/AKT pathway are critical for invasive properties and malignant transformation in CRC (5). The two most common genetic PI3K/AKT pathway alterations in CRC are activating mutations in the PI3K catalytic subunit gene *PIK3CA* or loss of the pathway suppressor PTEN (6). PTEN loss occurs in approximately 30% of CRCs and has been functionally implicated in CRC proliferation and progression (7). Additionally, chronic inflammation is a major risk factor for CRC development and the PI3K/AKT pathway can also be activated by reactive oxygen species (ROS) generated by activated immune cells at inflammatory sites (8,9).

Polycomb repressive complex 2 (PRC2) is a chromatin regulator that is involved in human development, tissue homeostasis, and cancer (10,11). Core components of PRC2 include EZH2 (enhancer of zeste homolog 2), Suz12 (suppressor of zeste 12), and EED (embryonic ectoderm development) (10). EZH2 functions as a lysine methyltransferase and EZH2-containing PRC2 canonically catalyzes trimethylation of histone 3 at lysine 27 (H3K27me3), which leads to chromatin compaction and transcriptional repression (12). H3K27me3 is involved in establishing chromatin domains that maintain transcriptional repression of hundreds of lineage-specific genes, reinforcing the maintenance of the current gene expression program and upholding cell identity (13,14). Initial studies showed that EZH2 acts as an oncogene by promoting cancer proliferation and progression (15). However, other studies revealed tumor suppressive functions of EZH2, highlighting unanticipated complexity in the role of PRC2 in cancer (16). Additionally, whereas EZH2 was consistently reported to be overexpressed in colon cancers, EZH2 expression levels have been found to be correlated positively or negatively with the survival of CRC patients depending on the study (17,18). These conflicting results suggest that there are contextdependent functions of EZH2 in cancer.

Emerging studies suggest that EZH2 can act noncanonically to regulate gene expression. For example, EZH2 interacts with and methylates various transcription factors, including androgen receptor (AR), GATA Binding Protein 4 (GATA4), related orphan receptor A (RORa), signal transducer and activator of transcription 3 (STAT3) and β-catenin, which either positively or negatively regulates their transcriptional activity (19–23). For example, EZH2-mediated methylation of STAT3 and GATA4 enhances and represses, respectively, the transcriptional activity of these transcription factors (19,22). Additionally, phosphorylation of EZH2 at Serine 21 (pS21-EZH2) by AKT has been shown to mediate the noncanonical roles of EZH2 in several of these studies. For example, AKT-mediated pS21-EZH2 induces EZH2 to interact with AR and STAT3 to induce the transcriptional activity of these transcription factors in prostate cancer and glioblastoma stem cells, respectively (20,22).

Here, we investigated the non-canonical role of EZH2 in CRC. We show that, following activation of the PI3K/AKT pathway, AKT phosphorylates EZH2. We further demonstrate that phosphorylation of EZH2 by active AKT is required for EZH2 to interact with and methylate β-catenin, a key transcription factor in CRC. Methylation of β-catenin regulates its transcriptional activity such that it has increased genome-wide chromatin binding and increased interaction with T-cell factor 1 (TCF1) and RNA polymerase II (RNAPII). Our work suggests that following phosphorylation by AKT, EZH2 fine-tunes β-catenin’s transcriptional activity resulting in modulation of the expression of genes involved in cell migration, and metabolic processes.

## Results

### AKT activation mediates EZH2 phosphorylation at Serine 21

Because AKT has previously been demonstrated to regulate EZH2 in other cancer types (20,22) and the PI3K/AKT pathway is commonly activated in CRC (7), we explored connections between AKT activity and EZH2 in CRC. To experimentally determine if AKT regulates EZH2 in CRC, we activated AKT by H_2_O_2_ treatment, which resulted in increased interaction between AKT and EZH2 in SW480 and RKO CRC cell lines and CRC organoids (Figure 1A, S1A, 1B). As a negative control, EZH2 was not detected in AKT IPs from H_2_O_2_-treated EZH2 knockdown (KD) cells and EZH2 KD did not alter the H_2_O_2_-induced increase in whole cell levels of AKT phosphorylated on serine 473 (pS473-AKT) (Figure 1A). AKT phosphorylates EZH2 at Serine 21 (S21) to regulate EZH2’s activity in prostate cancer and glioblastoma stem cells (20,22). To test if EZH2 is phosphorylated by AKT in CRC, we utilized an anti-pAKT substrate antibody, which recognizes the phosphorylated consensus sequence in AKT substrates (RXXS*/T*). Anti-pAKT substrate signal at EZH2’s expected size was increased in EZH2 IPs from H_2_O_2_-treated compared to untreated SW480 cells and CRC organoids (Figure 1C, S1B). The H_2_O_2_-induced increase in the EZH2 anti-pAKT substrate signal was blocked when cells were co-treated with AKT inhibitor (GSK690693) and H_2_O_2_, suggesting that the increase in phosphorylated EZH2 following H_2_O_2_ is caused by AKT activity (Figure 1C). AKT inhibitor treatment did not alter EZH2 levels in the nuclear lysates used for the IPs (input; Figure 1C). To determine if EZH2 phosphorylation is occurring at S21, we performed anti-HA IPs in SW480 cells transfected with HA-tagged EZH2 wild type (WT) and HA-tagged EZH2 S21A phospho-null (PN) plasmids. The EZH2 pAKT substrate band was detected in HA-EZH2 WT but not in HA-EZH2 S21A immunoprecipitates from cells treated with H_2_O_2_ (Figure 1D). Altogether, these results suggest that AKT activation in CRC induces AKT to interact with and phosphorylate EZH2 at S21.

**Figure 1.**
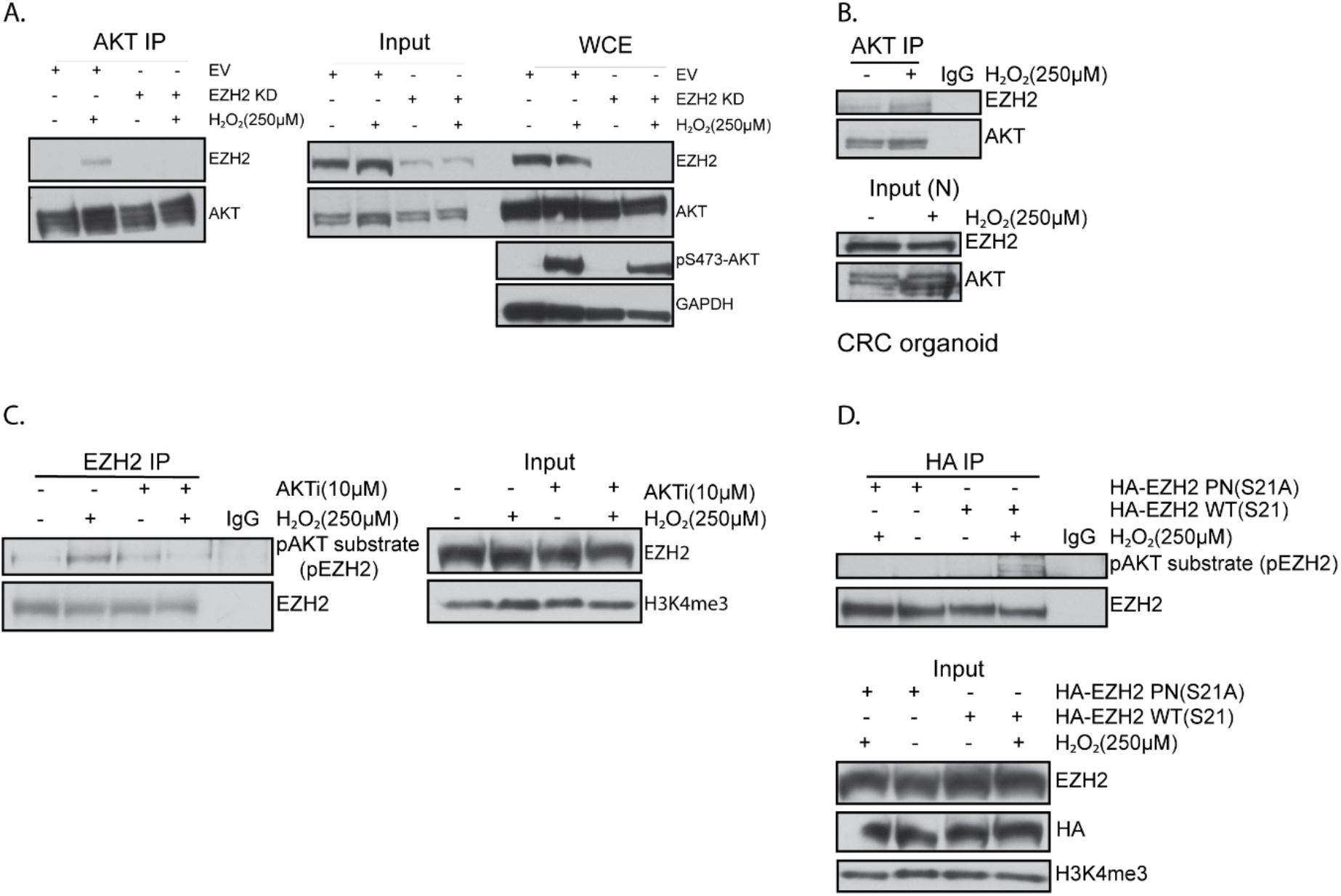
AKT activation mediates EZH2 phosphorylation at Serine 21. **A**. Western blots of AKT immunoprecipitations (IP) performed using nuclear lysates prepared from EZH2 knockdown (KD) and empty vector (EV) SW480 cells untreated or treated with 250 μM H_2_O_2_ for 30 minutes. Input is nuclear lysates used for IP and whole cell extract (WCE) is total protein prior to cellular fractionation. **B**. AKT IP performed using nuclear lysates prepared from human colorectal cancer organoids treated as in A. IP with IgG serves as a negative control. **C**. EZH2 IP performed using nuclear lysates prepared from SW480 cells untreated or treated with 10 μM AKT inhibitor (AKTi, GSK-690693) for 48 hours followed by no additional treatment or co-treatment with 250 μM H_2_O_2_ for 30 minutes. **D**. HA IP performed using nuclear lysates prepared from HA-EZH2 wildtype (WT) and HA-EZH2 phospho-null (PN) S21A expressing SW480 cells treated as in A.

### AKT-mediated EZH2 phosphorylation at S21 induces EZH2 to interact with RNAPII

EZH2 phosphorylation at S21 induces EZH2 to act as a transcriptional coactivator in other cancer types (20,22). To preliminarily determine if AKT activation switches EZH2’s function to become noncanonically involved in transcriptional activation, we tested if EZH2 interacts with RPB1, the largest subunit of the RNAPII complex. Treating SW480 cells with H_2_O_2_ induced EZH2 to interact with RNAPII without altering RNAPII or EZH2 protein levels in the whole cell or nuclear input fractions (Figure 2A, 2B). Interestingly, inhibiting AKT activity abolished the H_2_O_2_-induced increase in interaction between EZH2 and RNAPII (Figure 2B). As a control for the effectiveness of AKT inhibition, AKT inhibitor-treated samples had decreased levels of phosphorylated GSK3β, a known substrate for AKT, in input and whole cell fractions as compared to non-inhibitor-treated samples (Figure 2B). For the IPs in Figures 2A &B, RNaseA was added to the nuclear lysates to exclude the possibility that EZH2 was interacting with RNAPII through RNA bridging as has been shown previously (24,25). Additionally, H_2_O_2_ increased the interaction of AKT with RNAPII (Figure 2C). However, EZH2 KD decreased the H_2_O_2_-induced AKT-RNAPII interaction without altering total RNAPII or AKT levels in the nuclear input fractions (Figure 2C). Finally, to determine if EZH2 phosphorylation at S21 drives EZH2 to interact with RNAPII, HA IPs were performed in SW480 cells expressing similar levels of HA-EZH2 WT or HA-EZH2 S21A. Intriguingly, RNAPII was detected in HA-EZH2 WT but not in HA-EZH2 S21A immunoprecipitates from cells treated with H_2_O_2_ (Figure 2D). Altogether, these data suggest that phosphorylation of EZH2 at S21 via AKT induces EZH2 to interact with RNAPII.

**Figure 2.**
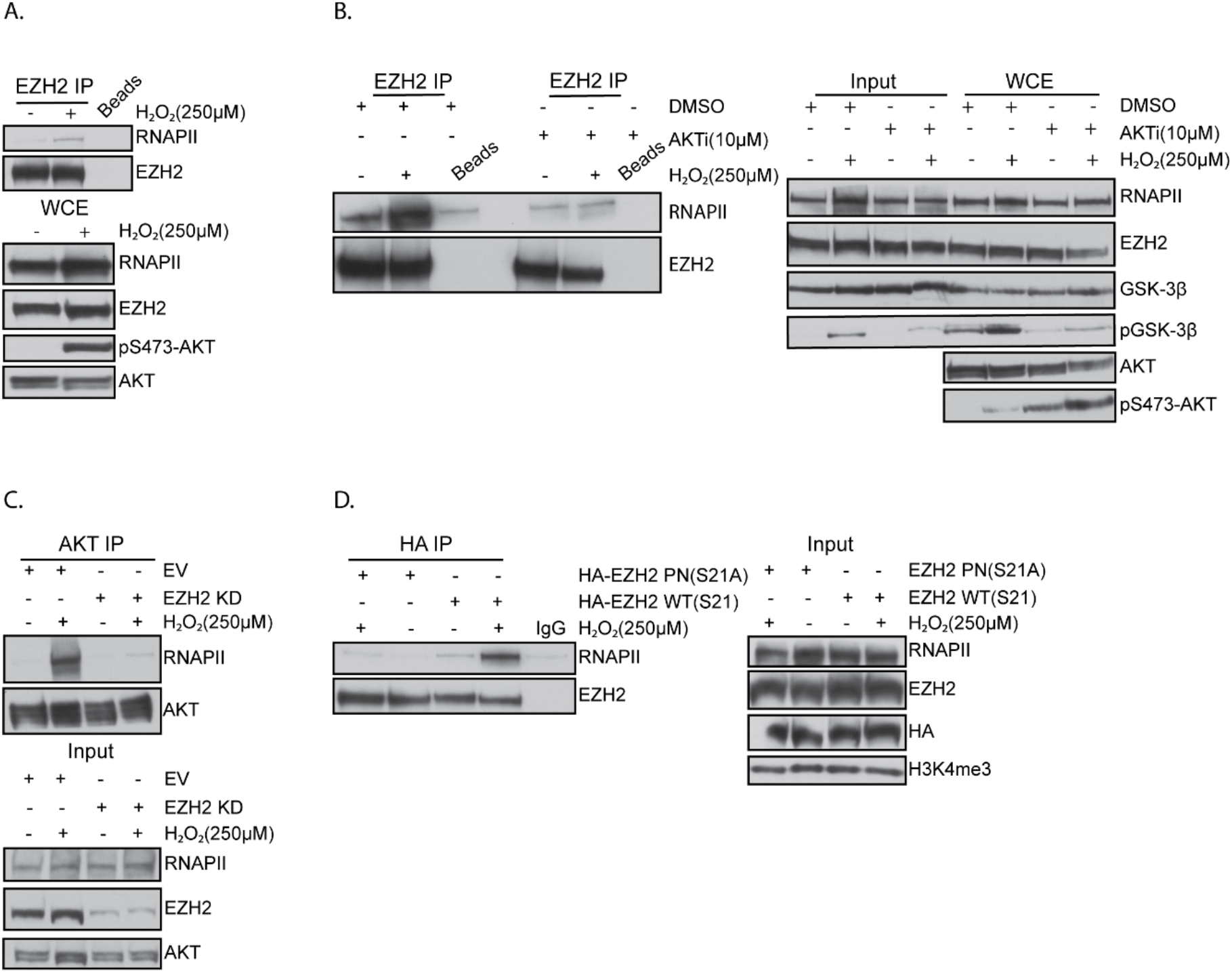
AKT-mediated EZH2 phosphorylation at S21 induces EZH2 to interact with RNAPII. **A**. Western blots of EZH2 immunoprecipitations (IP) performed using nuclear lysates prepared from SW480 cells untreated or treated with 250 μM H_2_O_2_ for 30 minutes. IP with beads serves as a negative control. Whole cell extract (WCE) is total protein prior to cellular fractionation. **B**. EZH2 IP performed using nuclear lysates prepared from SW480 cells treated with 10 μM AKT inhibitor (AKTi, GSK-690693) or DMSO for 48 hours followed by no additional treatment or cotreatment with 250 μM H_2_O_2_ for 30 minutes. IP with Beads serves as a negative control. Input is nuclear lysates used for IP. **C**. AKT IP performed using nuclear lysates prepared from EZH2 knockdown (KD) and empty vector (EV) SW480 cells treated as in A. **D**. HA IP performed using nuclear lysates prepared from HA-EZH2 wildtype (WT) and HA-EZH2 phospho-null (PN) S21A expressing SW480 cells treated as in A. IP with IgG serves as a negative control.

### AKT-mediated EZH2 phosphorylation induces EZH2’s interaction with β-catenin

In prostate cancer, EZH2 phosphorylation at S21 promotes its interaction with AR to induce AR’s transcriptional activity (20). Because the WNT/β-catenin pathway is hyperactive in almost 90% of CRCs (4), we tested whether AKT-induced phosphorylation of EZH2 induced EZH2 to interact with β-catenin and if EZH2 mediated β-catenin’s interaction with RNAPII in CRC. H_2_O_2_ treatment increased the interaction between β-catenin and EZH2 in SW480 cells, CRC organoids, and HEK293T cells (Figure 3A, 3B, S2A). H_2_O_2_ did not alter β-catenin levels in the nuclear input fractions prepared from SW480 or HEK293T (Figure 3A, S2A). However, β-catenin levels were increased in the nuclear input fraction prepared from CRC organoid cells following H_2_O_2_ treatment (Figure 3B). PTEN loss, which activates the PI3K/AKT pathway, is also common in CRC (7). Interestingly, we identified a significant negative correlation between PTEN and EZH2 expression in TCGA CRC samples (Figure S2B). As expected, PTEN KD in SW480 cells induced AKT phosphorylation (Figure S2C). Therefore, we performed additional experiments in PTEN KD cells to complement our findings using H_2_O_2_ treatment. Like H_2_O_2_ treatment, PTEN KD in SW480 cells increased the interaction between EZH2 and RNAPII compared to empty vector (EV) KD cells (Figure S2D).

**Figure 3.**
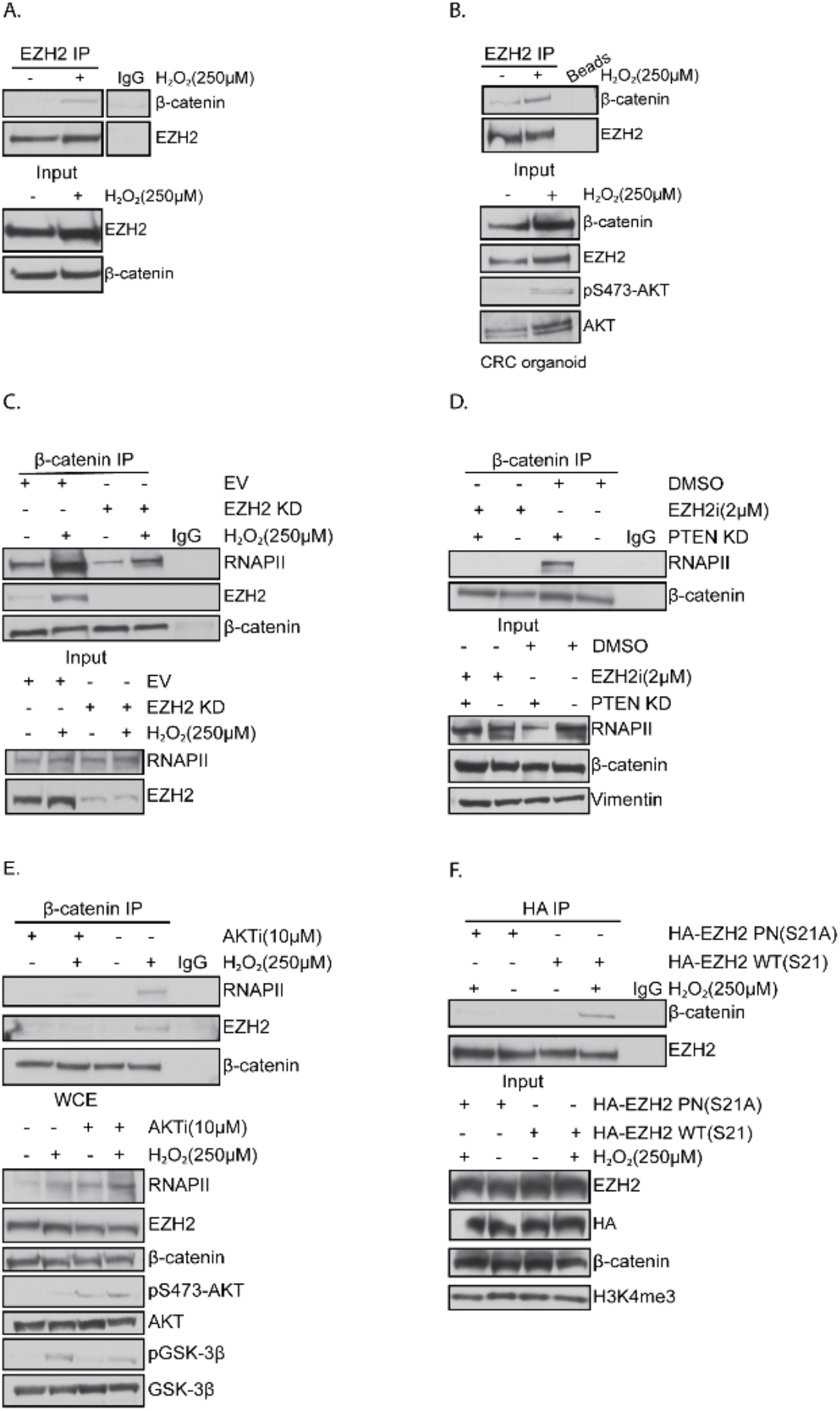
AKT-mediated EZH2 phosphorylation induces EZH2’s interaction with β-catenin. **A**. Western blots of EZH2 immunoprecipitations (IP) performed using nuclear lysates prepared from SW480 cells untreated or treated with 250 μM H_2_O_2_ for 30 minutes. IgG IP serves as a negative control. Input is nuclear lysates used for IP. **B**. EZH2 IP performed using nuclear lysates prepared from human colorectal cancer organoids treated as in A. **C**. β-catenin IP performed using nuclear lysates prepared from EZH2 knockdown (KD) and empty vector (EV) SW480 cells treated as in A. **D**. β-catenin IP performed using nuclear lysates prepared from PTEN KD and EV SW480 cells treated with 2 μM EZH2 inhibitor (EZH2i, GSK-503) or DMSO for 72 hours. **E**. β-catenin IP preformed using nuclear lysates prepared from SW480 cells untreated or treated with 10 μM AKT inhibitor (AKTi, GSK-690693) for 48 hours followed by no additional treatment or co-treatment with 250 μM H_2_O_2_ for 30 minutes. Whole cell extract (WCE) is total protein prior to cellular fractionation. **F**. HA IP performed using nuclear lysates prepared from HA-EZH2 wildtype (WT) and HA-EZH2 phospho-null (PN) S21A expressing SW480 cells treated as in A.

Based on our findings so far, we hypothesized that EZH2 may regulate β-catenin’s interaction with the transcriptional machinery. H_2_O_2_ and PTEN KD induced an increase in interaction between β-catenin and RNAPII compared to untreated cells (Figure S2E). To determine if EZH2 played a role in this interaction, we treated EV and EZH2 KD SW480 cells with H_2_O_2_, which increased β-catenin’s interaction with RNAPII and EZH2 in EV cells (Figure 3C). EZH2 KD drastically reduced the H_2_O_2_-induced interaction between β-catenin and RNAPII without altering RNAPII levels in the nuclear input fraction (Figure 3C). As a control, EZH2 was not detected in β-catenin IPs in EZH2 KD cells (Figure 3C). Furthermore, treating SW480 cells with an EZH2 inhibitor (GSK503, EZH2i) blocked the PTEN KD-mediated β-catenin-RNAPII interaction (Figure 3D). We hypothesized that AKT activation is required for EZH2 to interact with β-catenin and RNAPII. Consistent with this hypothesis, inhibiting AKT activity reduced the interaction of β-catenin with RNAPII and EZH2 after H_2_O_2_ treatment without affecting the levels of EZH2, RNAPII, or β-catenin in the whole cell extract fraction (Figure 3E). To test whether EZH2 phosphorylation at S21, which is mediated by AKT, drives EZH2 to interact with β-catenin, we transfected SW480 cells with HA-tagged WT and phosphorylation null EZH2 constructs and performed anti-HA IPs. Interestingly, H_2_O_2_ treatment and PTEN KD induced HA-EZH2 WT but not HA-EZH2 S21A to interact with β-catenin (Figure 3F, S3). These data suggest that phosphorylation of EZH2 at S21 via active AKT induces EZH2 to interact with RNAPII and β-catenin. In addition, β-catenin’s interaction with RNAPII likely requires EZH2 enzymatic activity because inhibiting EZH2 activity blocked the AKT activity-dependent interaction between β-catenin and RNAPII.

### EZH2 mediates β-catenin trimethylation at K49

EZH2 has been reported to methylate non-histone proteins, including β-catenin (19,21–23), and we have determined that AKT activation induced EZH2 to interact with β-catenin. To test if H_2_O_2_ induces β-catenin methylation, we utilized FLAG-tagged constructs of WT β-catenin or β-catenin mutants that are methylation null at one or both of the reported sites of β-catenin methylation, K19 and K49 (26). H_2_O_2_ treatment increased lysine trimethylation of immunoprecipitated FLAG-β-catenin WT and FLAG-β-catenin K19R relative to FLAG-β-catenin WT immunoprecipitated from untreated cells (Figure 4A). However, the level of lysine trimethylation of FLAG-β-catenin K49R and FLAG-β-catenin K14/49R immunoprecipitated from H_2_O_2_-treated cells was not increased (Figure 4A). There were no differences in mono-methyl lysine signal among the different samples and all samples expressed similar levels of FLAG-β-catenin in the nuclear input fraction (Figure 4A). The H_2_O_2_-induced increase in trimethylation of FLAG-β-catenin WT but not FLAG-β-catenin K49R was confirmed in a second experiment (Figure S4). The reverse IPs using an anti-trimethyl lysine antibody showed similar results with the FLAG signal at the expected size of β-catenin being increased in immunoprecipitates from H_2_O_2_ treated cells expressing FLAG-β-catenin WT but not FLAG-β-catenin K49R relative to their respective untreated controls (Figure 4B). PTEN KD also increased trimethylation of FLAG-β-catenin WT, not FLAG-β-catenin K49R relative to their respective EV controls (Figure 4C). These results suggested that the trimethylation of β-catenin was occurring at K49 in response to AKT activation. Our finding that AKT activation induced EZH2 to interact with β-catenin (Figure 2) led us to hypothesize that β-catenin trimethylation is mediated by EZH2. FLAG IPs in SW480 cells transfected with FLAG-β-catenin WT and β-catenin IP in EV and PTEN KD cells confirmed that H_2_O_2_ and PTEN KD increased β-catenin trimethylation (Figure 4D, 4E). Consistent with our hypothesis, EZH2 inhibition reduced H_2_O_2_ and PTEN KD-mediated β-catenin trimethylation without altering total levels of FLAG-β-catenin or β-catenin (Figure 4D, 4E). These results suggest that activation of AKT induces EZH2 to interact with and trimethylate β-catenin at K49.

**Figure 4.**
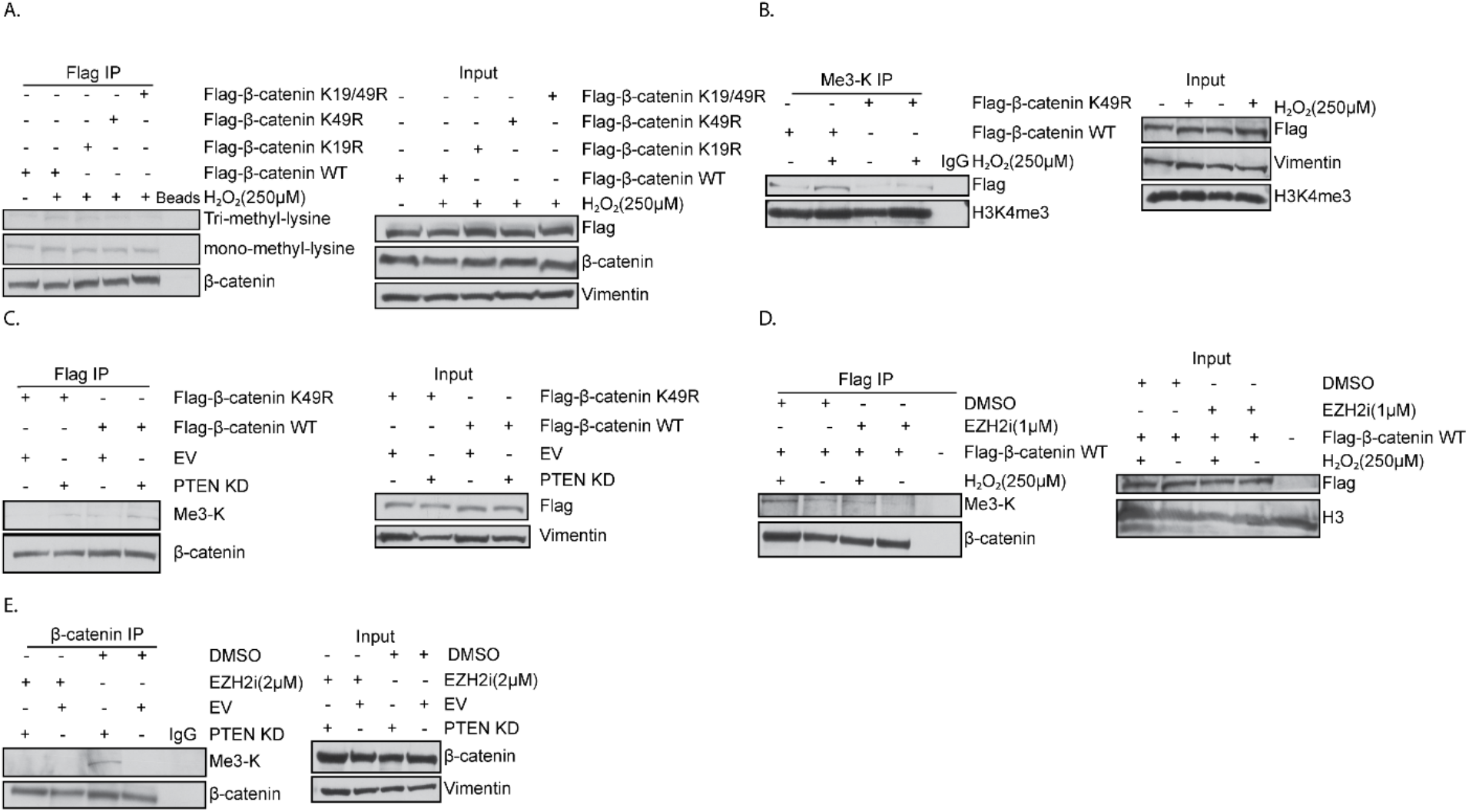
EZH2 mediates β-catenin trimethylation at K49. **A**. Western blots of Flag immunoprecipitations (IP) performed using nuclear lysates prepared from Flag β-catenin wildtype (WT), Flag β-catenin K19R, Flag β-catenin K49R and Flag β-catenin K19/49R expressing SW480 cells treated with 250 μM H_2_O_2_ for 30 minutes and Flag β-catenin WT expressing untreated cells. Input is nuclear lysates used for IP. IP with beads only serves as a negative control. **B**. Trimethyl lysine (Me3-K) IP performed using nuclear lysates prepared from Flag β-catenin WT and Flag β-catenin K49R expressing SW480 cells treated as in A. IgG IP serves as a negative control. **C.** Flag IP performed using nuclear lysates prepared from Flag-tagged β-catenin WT and Flag-tagged β-catenin K49R expressing PTEN KD or empty vector (EV) SW480 cells. **D**. Flag IP performed using nuclear lysates prepared from Flag-tagged β-catenin WT expressing SW480 cells treated with 2 μM EZH2 inhibitor (EZH2i, GSK-503) or DMSO for 72 hours and with or without co-treatment with 250 μM H_2_O_2_ for 30 minutes. Cells not expressing Flag plasmid were used as a negative control for the IP. **E**. β-catenin IP preformed using nuclear lysates prepared from EV or PTEN KD SW480 cells treated with EZH2 inhibitor as in D.

### EZH2-mediated trimethylation of β-catenin induces β-catenin to interact with TCF1 and RNAPII

During WNT pathway activation, β-catenin interacts with TCF proteins to induce WNT target gene expression (27). TCF1 IPs in SW480 cells showed that the interaction between TCF1 and β-catenin slightly increased after H_2_O_2_ treatment (Figure 5A, S5A). EZH2 inhibition or KD in SW480 cells reduced the interaction of TCF1 with β-catenin and RNAPII (Figure 5A, S5A). The reverse IPs of FLAG-β-catenin WT showed a more robust H_2_O_2_-induced interaction between β-catenin and TCF1 and treating cells with EZH2 inhibitor decreased this interaction (Figure 5B). β-catenin and TCF1 IPs demonstrated that PTEN KD also induced β-catenin to interact with TCF1 and inhibiting EZH2 activity reduced this interaction as well (Figure 5C, S5B). Because EZH2 inhibition reduced the interaction of β-catenin and TCF1, we hypothesized that this interaction may be promoted by EZH2 trimethylating β-catenin. To test this hypothesis, FLAG IPs were performed in SW480 cells expressing similar levels of FLAG-β-catenin WT and FLAG-β-catenin K49R. H_2_O_2_ treatment increased the interaction of FLAG-β-catenin WT but not FLAG-β-catenin K49R with TCF1 (Figure 5D). PTEN KD slightly increased the interaction of FLAG-β-catenin WT with TCF1 relative to EV whereas the interaction between FLAG-β-catenin K49R and TCF1 was reduced relative to FLAG-β-catenin WT in EV and PTEN KD cells (Figure 5E). Additionally, FLAG-β-catenin K49R had reduced interaction with RNAPII than FLAG-β-catenin WT in PTEN KD SW480 cells (Figure 5E). Altogether, these findings suggest that EZH2-mediated trimethylation of β-catenin induces β-catenin’s interaction with RNAPII and TCF1 after AKT activation.

**Figure 5.**
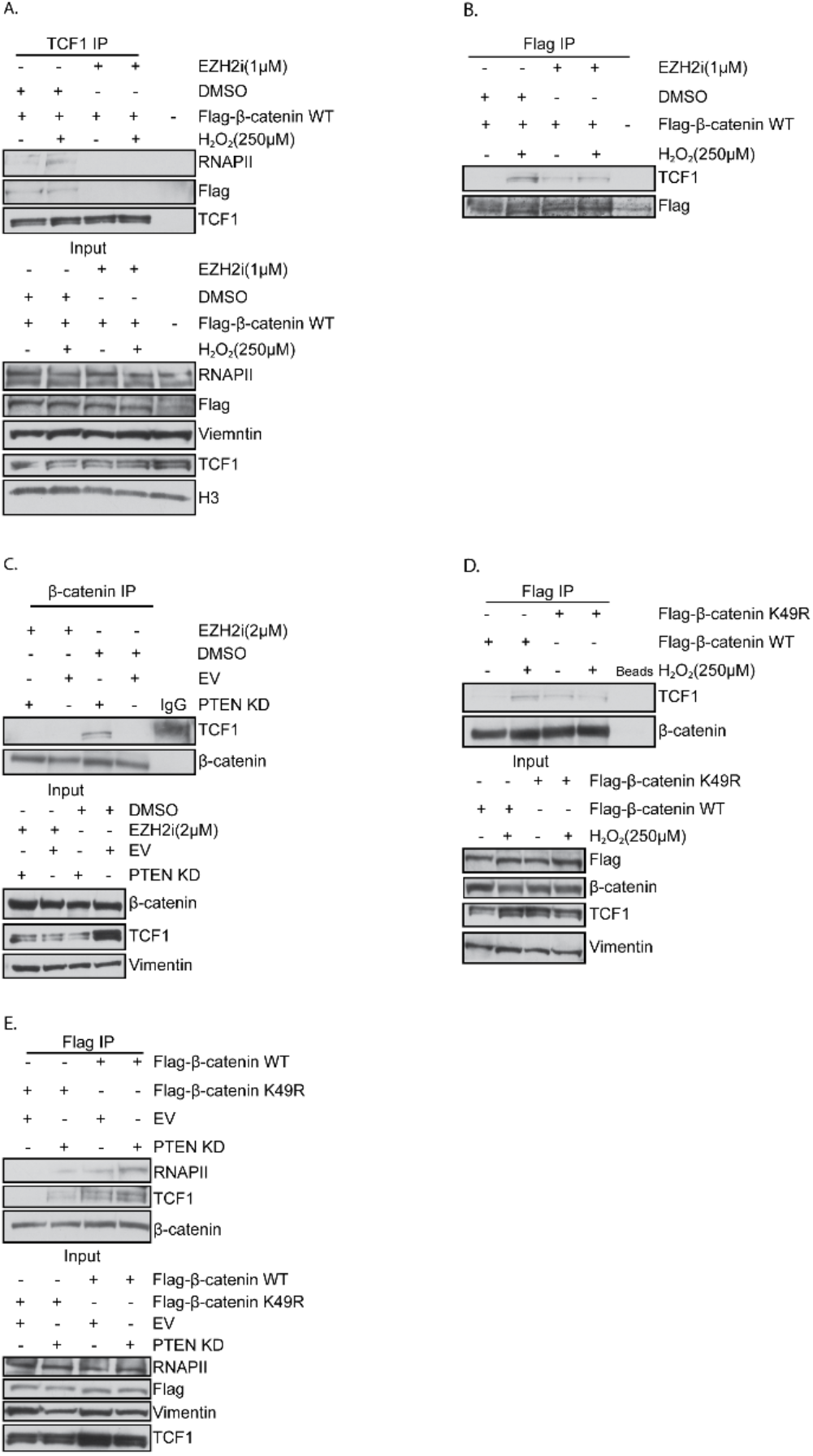
EZH2-mediated trimethylation of β-catenin induces β-catenin to interact with TCF1 and RNAPII. **A**. Western blots of TCF1 immunoprecipitation (IP) performed using nuclear lysates prepared from Flag-tagged β-catenin wildtype (WT) expressing SW480 cells treated with 1 μM EZH2 inhibitor (EZH2i, GSK-503) or DMSO for 72 hours with or without co-treatment with 250 μM H_2_O_2_ for 30 minutes. IgG IP serves as a negative control. Input is nuclear lysates used for IP. **B**. Flag IP performed using nuclear lysates prepared from Flag-tagged β-catenin WT expressing SW480 cells treated as in A. IPs were performed using the same nuclear lysate in A. **C**. β-catenin IP performed using nuclear lysates prepared from PTEN knockdown (KD) or empty vector (EV) SW480 cells treated with 2 μM EZH2 inhibitor (EZH2i, GSK-503) or DMSO for 72 hours. **D**. Flag IP performed using nuclear lysates prepared from Flag-tagged β-catenin WT and Flag-tagged β-catenin K49R expressing SW480 cells untreated or treated with 250 μM H_2_O_2_ for 30 minutes. IP with beads only serves as a negative control. **E**. Flag IP performed using nuclear lysates prepared from Flag-tagged β-catenin WT and Flag-tagged β-catenin K49R expressing PTEN KD or EV SW480 cells.

To confirm our hypothesis that AKT activation induces AKT to interact with and phosphorylate EZH2, we induced AKT activation by performing base editing of the *PIK3CA* genomic loci. Glutamic acid was replaced by lysine at 453 or 545 (E453K or E545K) (Figure S5C) producing common gain of function mutations of *PIK3CA* that resulted in increased pAKT levels (Figure S5D). SW480 cells harboring the *PIK3CA* E453K or E545K mutation had increased EZH2 interaction with and phosphorylation by AKT relative to control EV SW480 cells with WT *PIK3CA* (Figure S5E). *PIK3CA* E453K and E545K cells also had increased interaction of β-catenin with RNAPII, EZH2, and TCF1 relative to EV control cells (Figure S5F).

### EZH2 phosphorylation increases its binding to chromatin

Next, we wanted to determine how the AKT-EZH2-β-catenin axis altered the binding of these proteins to chromatin. Because we demonstrated that phosphorylation of EZH2 at S21 by AKT caused EZH2 to interact with RNAPII and β-catenin (Figure 2D, 3F), we tested whether EZH2 phosphorylation affects EZH2’s binding to chromatin. EZH2 KD SW480 cells were rescued with HA-EZH2 WT, HA-EZH2 S21A, or EV followed by H_2_O_2_ treatment. H_2_O_2_ treatment increased EZH2 levels relative to mock treatment in the chromatin fraction isolated from HA-EZH2 WT-expressing cells (Figure 6A). Chromatin-bound EZH2 was not detected in cells expressing either HA-EZH2 S21A or EV even though levels of EZH2 in the whole cell fraction were similar between HA-EZH2 WT and HA-EZH2 S21A expressing cells (Figure 6A). Additionally, EZH2 levels were higher in the chromatin fraction of cells rescued with an EZH2 phosphomimetic mutant construct, HA-EZH2 S21D, relative to cells rescued with HA-EZH2 WT or EV (Figure 6B). EZH2 KD SW480 cells expressing HA-EZH2 WT also had increased levels of chromatin-bound TCF1 after H_2_O_2_ treatment than mock-treated or HA-EZH2 S21A expressing cells (Figure 6A). We next tested if the binding of RNAPII and/or TCF1 to chromatin is dependent on AKT and/or EZH2 catalytic activity as we have shown the activity of these enzymes is required for β-catenin to interact with RNAPII and TCF1. Treating cells with AKT or EZH2 inhibitors blocked the H_2_O_2_-induced increase of RNAPII and TCF1 levels in the chromatin fraction but had minimal effect on the chromatin levels of these proteins in untreated cells (Figure 6C). While AKT and EZH2 inhibitors did not alter the levels of RNAPII in whole cell fractions, they reduced the total level of TCF1 (Figure 6C). Because we have shown that the interaction of β-catenin with TCF1 and RNAPII is dependent on β-catenin‘s methylation by EZH2 (Figure 3D, 5), we tested if trimethylation of β-catenin alters β-catenin’s chromatin binding. Interestingly, PTEN KD induced an increase in FLAG-β-catenin WT but not FLAG-β-catenin K49R levels in chromatin fraction (Figure 6D). Altogether, these results suggest that phosphorylation of EZH2 by AKT regulates EZH2’s binding to chromatin. Additionally, EZH2-mediated β-catenin trimethylation regulates β-catenin’s binding to chromatin.

**Figure 6.**
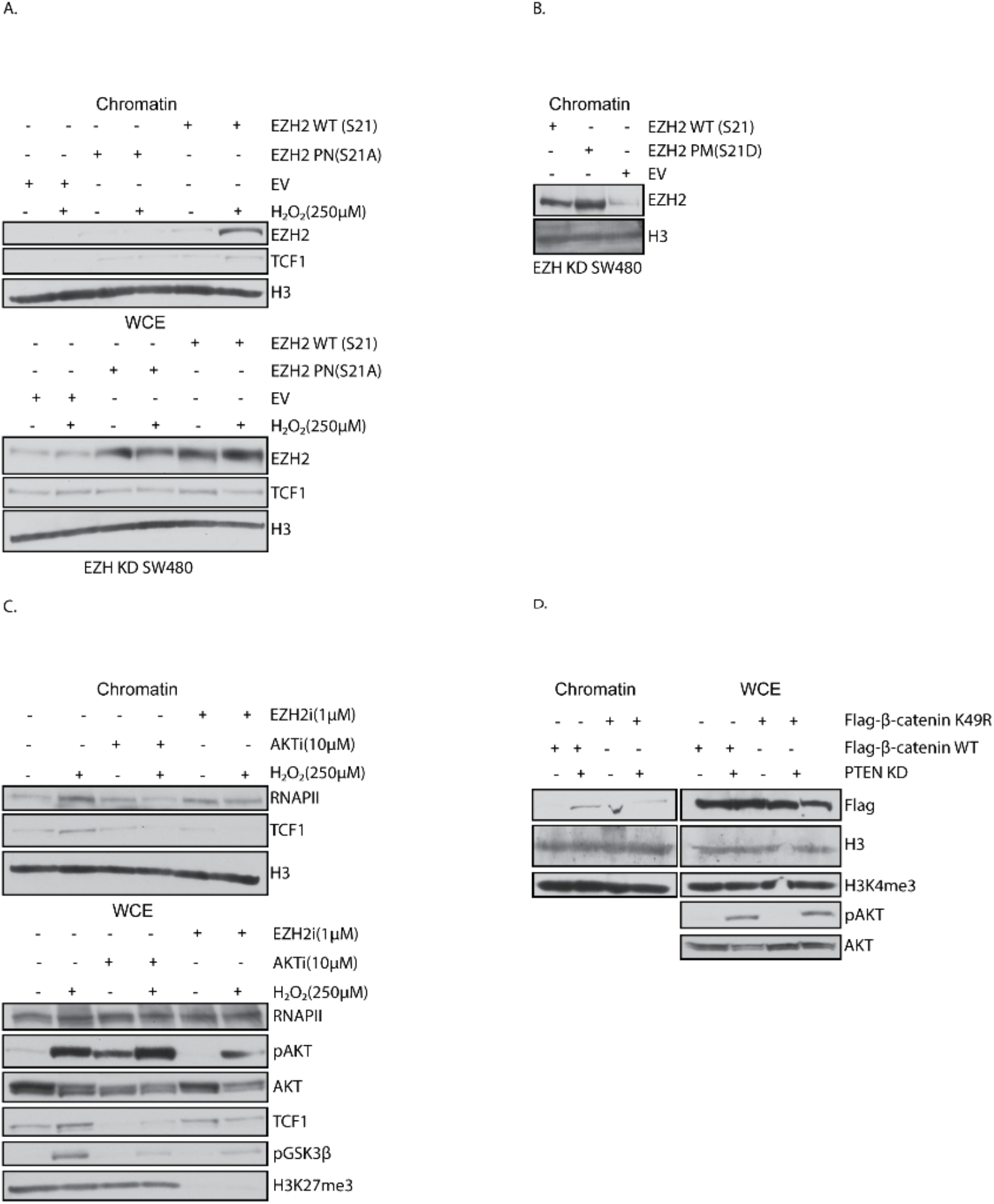
Phosphorylation of EZH2 increases its binding to chromatin. **A**. Western blots of chromatin lysate prepared from HA-EZH2 wildtype (WT), HA-EZH2 phospho-null (PN) S21A and HA-empty vector (EV) expressing SW480 cells treated with 250 μM H_2_O_2_ for 30 minutes. Whole-cell extract (WCE) serves as a control for the chromatin extract. **B**. Western blots of chromatin lysate prepared from HA-EZH2 WT, HA-EZH2 S21D and HA-EV expressing SW480 cells. **C**. Western blots of chromatin lysate prepared from SW480 cells untreated and treated with 1 μM EZH2 inhibitor (EZH2i, GSK-503) for 72 hours or with 10 μM AKT inhibitor (AKTi, GSK-690693) for 48 hours and followed by no additional treatment or co-treatment with 250 μM H_2_O_2_ for 30 minutes. **D**. Western blots of chromatin lysate prepared from FLAG-β-catenin WT and FLAG-β-catenin K49R expressing PTEN knockdown (KD) and EV expressing SW480 cells. WCE serves as a control for the chromatin extract.

### EZH2 inhibition blocks the PTEN knockdown-induced increase in β-catenin enrichment over the genome

We previously demonstrated that phosphorylation of EZH2 at S21 increased EZH2’s binding to chromatin (Figure 6A). To determine how phosphorylation alters the genome-wide binding pattern of EZH2, CUT&RUN for HA-tagged EZH2 in PTEN KD and EV SW480 cells transfected with HA-EZH2 WT or HA-EZH2 S21A was performed. A portion of the called HA-EZH2 WT peaks in PTEN KD overlapped with the called peaks in EV cells (84 peaks; Figure 7A, top). A larger portion of the called HA-EZH2 WT peaks were specific for PTEN KD SW480 cells (942 peaks; Figure 7A, bottom). There were only 64 HA-EZH2 WT peaks uniquely called in EV cells. Cells transfected with HA-EZH2 S21A had a very low number of called peaks in EV (40 peaks) and PTEN KD (41 peaks) samples, the majority of which were also present in the HA-EZH2 WT samples (Figure 7A). This data confirmed our hypothesis that phosphorylation of EZH2 by AKT increases EZH2 binding to chromatin. Our chromatin data suggested that EZH2-mediated β-catenin trimethylation increased β-catenin binding to the chromatin (Figure 6D). To systematically define the genomewide enrichment pattern of β-catenin, ChIP-seq for FLAG-tagged β-catenin was performed in EV and PTEN KD SW480 cells. PTEN KD increased the enrichment of FLAG-β-catenin over the chromatin in an EZH2 activity-dependent manner (Figure 7C, 6SA). Similar to HA-tagged EZH2, a portion of the called FLAG-β-catenin peaks in PTEN KD cells overlapped with FLAG-tagged β-catenin peaks in EV cells (2525 peaks; Figure 7B, middle). A larger portion of the called FLAG-tagged β-catenin peaks were specific only for PTEN KD cells (8453 peaks; Figure 7B, bottom). There were an additional 2323 FLAG-tagged β-catenin peaks uniquely called in EV cells (Figure 7B, top). Inhibiting EZH2 drastically reduced the number of called FLAG-β-catenin peaks in EV and PTEN KD cells to 58 and 288 peaks, respectively, all of which were also present in the noninhibitor treated samples (Figure 7B). Profile plots of the PTEN KD-specific peaks further demonstrated that FLAG-tagged β-catenin enrichment at these peaks decreased in EV cells and was further decreased in EZH2 inhibitor-treated EV and PTEN KD cells (Figure 7C). This finding was consistent in a second ChIP-seq experiment (Figure S6A). Altogether this data suggests that EZH2 phosphorylation and subsequent β-catenin trimethylation increase the enrichment of EZH2 and β-catenin at numerous loci across the genome.

**Figure 7.**
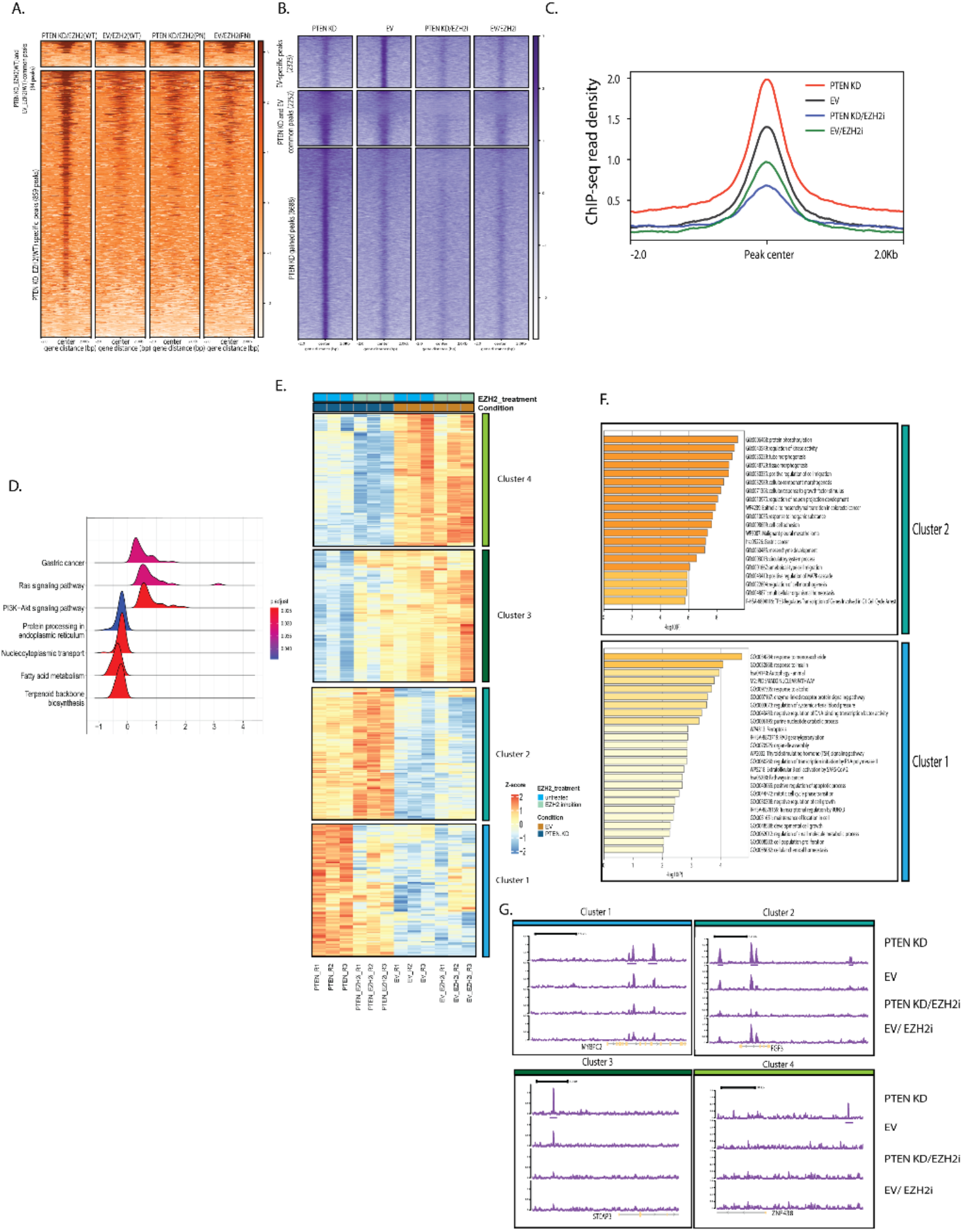
EZH2 inhibition blocks the PTEN knockdown-induced increase in β-catenin enrichment over the genome. **A**. Metagenomic heatmap for HA CUT&RUN prepared from HA-EZH2 wildtype (WT) and HA-EZH2 S21A expressing PTEN knockdown (KD) and empty vector (EV) SW480 cells. **B**. Metagenomic heatmap for FLAG ChIP-seq prepared from FLAG-β-catenin WT and FLAG-β-catenin K49R expressing PTEN KD and EV expressing SW480 cells. **C.** Average ChIP-seq read intensity for all peaks shown in B. **D**. Ridgeplot for differentially expressed genes (DEGs) in PTEN KD vs EV SW480 cells. **E**. Heatmap for DEGs in PTEN KD versus EV clustered manually based on the effect of EZH2 inhibitor (GSK503, 1 μM, 72 hours). **F.** Functional gene annotations for DEGs in cluster 1 and cluster 2 from E generated by Metascape. **G**. ChIP-seq gene tracks of representative DEGs in PTEN KD versus EV SW480 cells.

### Increased enrichment of β-catenin across the genome regulates gene expression

To determine the impact of the AKT-EZH2-β-catenin axis on transcription, we performed RNA-seq with and without EZH2 inhibition in EV and PTEN KD SW480 cells. PTEN KD resulted in 265 up-regulated and 141 down-regulated genes compared to EV (|log 2FC|>0.5; P<0.05; Figure S6B). As expected, PTEN KD upregulated the expression of genes involved in the PI3K-AKT pathway whereas PTEN KD down-regulated genes were involved in metabolic processes such as fatty acid metabolism (Figure 7D). Inhibiting EZH2 activity altered the expression of PTEN KD-regulated genes (Figure 7E). Therefore, we manually clustered PTEN KD-regulated genes into four clusters based on the effect of the EZH2 inhibitor on their expression. In cluster one, EZH2 inhibition reduced the expression of PTEN KD-upregulated genes (Figure 7E). GO analysis of genes in this cluster showed the enrichment of pathways related to signaling pathways and transcription initiation (Figure 7F). In cluster two, we detected a further increase in the expression of PTEN KD-upregulated genes in response to the EZH2 inhibitor (Figure 7E). GO analysis of this cluster indicated enrichment of pathways related to tissue morphogenesis, cell migration, and epithelial-to-mesenchymal transition (EMT) (Figure 7F). While the EZH2 inhibitor increased the expression of PTEN KD down-regulated genes in cluster three, it further reduced gene expression of PTEN KD down-regulated genes in cluster four (Figure 7E). GO analysis of these two clusters showed enrichment of pathways associated with different metabolic processes (Figure S6C).

Through the integration of genomic binding and RNA-seq profiles, we determined 25, 51, 14, and 11 genes from clusters 1, 2, 3, and 4, respectively, that exhibited increased binding of FLAG-β-catenin in PTEN KD cells (Figure S6D), as exemplified by *MYBPC2* from cluster 1 and *FGF3* from cluster 2 and *STEAP3* from cluster 3 and *ZNF438* from cluster 4 (Figure 7G). We also determined that 263 peaks had increased co-binding of both β-catenin and EZH2 in PTEN KD relative to EV cells such as *NTM* (Figure S6E). Additionally, a small number of EZH2 peaks overlapped with differentially expressed genes in response to PTEN KD as exemplified by *P2RX5* and *CLSTN2* (Figure S6F). However, the majority of EZH2 peaks gained in PTEN KD cells did not overlap with FLAG-β-catenin peaks and were not associated with differentially expressed genes in PTEN KD cells. Altogether these results suggest that EZH2-mediated β-catenin trimethylation fine-tunes the transcriptional activity of β-catenin to regulate gene expression in PTEN KD SW480 cells.

### EZH2 inhibition enhances cell migration in PTEN knockdown cells

GO analysis of cluster 2 genes that were defined as being increased in expression with PTEN KD and further increased with EZH2 inhibition demonstrated enrichment for genes involved in migration, morphogenesis, and epithelial to mesenchymal transition (EMT) (Figure 7F). Therefore, we decided to narrow our focus to cluster 2 genes and performed hallmark analysis using the Human MSigDB Collections (28). Consistent with our previous analysis, cluster 2 genes were significantly associated with apical junctions and EMT hallmarks (Figure 8A, B). β-catenin appeared to directly regulate some EMT-related genes as PTEN KD increased the binding of FLAG-β-catenin to their promoter regions, as exemplified by *ANO1* and *LAMA3* (Figure 8C). Because the cluster 2 genes were defined as being further increased by EZH2 inhibition we explored the effect of EZH2 inhibition alone on gene expression pathways using our RNA-seq data. Hallmark analysis of the 255 genes (log 2FC>0.5; P<0.05) up-regulated in response to EZH2 inhibitor in EV cells showed that the most significantly enriched hallmark was the EMT hallmark (Figure S7). This result is consistent with the known role of EZH2 in repressing EMT genes and maintaining cell identity (29–31). To determine how these changes in gene expression alter migration, we performed transwell migration assays. PTEN KD increased SW480 cell migration compared to EV (Figure 8D, E). While treatment of EV cells with EZH2 inhibitor trended to induce increased cell migration, EZH2 inhibitor treatment significantly increased PTEN KD cell migration. Our results suggest that PTEN KD increases expression of EMT-related genes and cell migration and treatment with EZH2 inhibitor enhances these PTEN KD-mediated effects.

**Figure 8.**
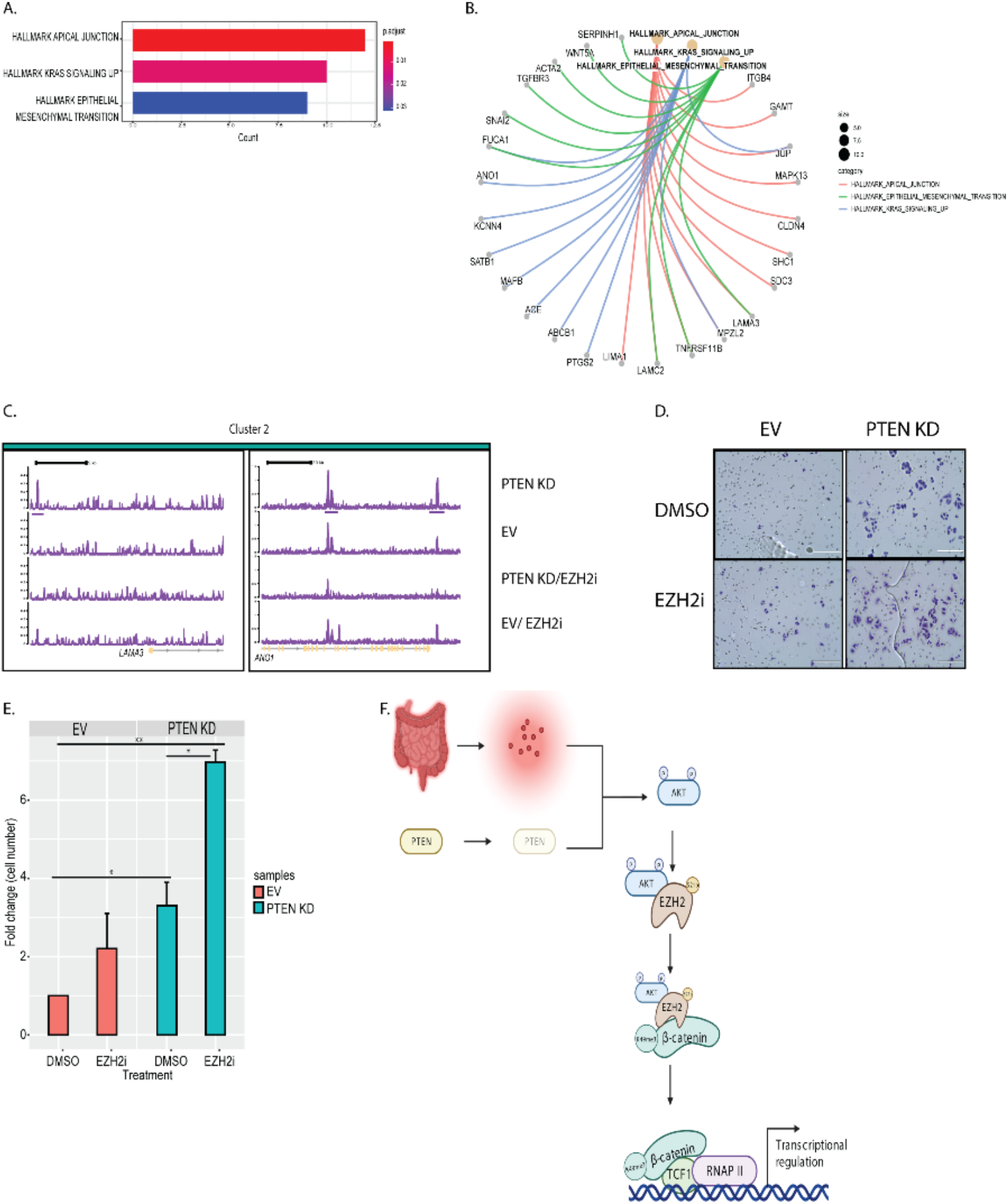
Increased enrichment of β-catenin across the genome regulates gene expression. **A**. Barplot for the hallmark analysis for differentially expressed genes (DEGs) in cluster 2. **B**. Centplot for the hallmark analysis in B. **C**. FLAG-β-catenin ChIP-seq gene tracks of representative hallmark genes. **D**. Empty vector (EV) and PTEN knockdown (KD) cells were treated with DMSO or 2 μM EZH2 inhibitor (EZH2i, GSK-503) for 72 hours followed by plating cells in the upper chamber of a transwell insert. Brightfield images of crystal violet–stained migrated cells were taken after 48 hours. Scale bar = 200 microns. **E**. Quantification of migration normalized to migration counts for untreated EV cells. Results are represented as mean +/- SD. Significance was determined by one-way ANOVA with the Tukey multiple comparisons test. All significant comparisons are shown. ** P< 0.01. **F**. Model depicting the role of pS21-EZH2 in regulating AKT-mediated transcription through β-catenin methylation.

## Discussion

In this study, we reveal the mechanism by which EZH2 regulates β-catenin in CRC. We determined that activation of PI3K/AKT, a pathway activated in a significant portion of CRCs, results in EZH2 phosphorylation at S21 (pS21-EZH2) in CRC, similar to what has been found in other cancer types (20,22). Our data further demonstrate that phosphorylation of EZH2 via AKT is the molecular switch that drives EZH2 to interact with and methylate β-catenin (Figure 8F). Inhibition of AKT signaling blocks EZH2’s interaction with and methylation of β-catenin. Additionally, a phospho-null EZH2 mutant (HA-EZH2 S21A) did not interact with β-catenin following AKT activation confirming that phosphorylation at S21 drives EZH2’s interaction with β-catenin. These findings agree with other studies in which phosphorylation of EZH2 by AKT induces EZH2’s interaction with other transcription factors such as AR and STAT3 in prostate cancer and glioblastoma, respectively (20,22).

Previously, EZH2 has been reported to modulate β-catenin’s transcriptional activity in other cell types. For example, in liver cancer stem cells, EZH2-mediated methylation of β-catenin at K49 increased β-catenin’s transcriptional activity and promoted WNT signaling, while in embryonic stem cells β-catenin methylation at the same site by EZH2 led to gene repression during development (23,32). Here, we demonstrated that EZH2 trimethylates β-catenin, which fine-tunes β-catenin’s transcriptional activity in CRC. We determined that EZH2’s catalytic activity is essential for modulating β-catenin regulated gene expression. Treating cells with an EZH2 inhibitor (GSK503) blocked the H_2_O_2_ or PTEN KD-mediated β-catenin interaction with RNAPII and TCF1. Additionally, a methylation-null mutant construct, FLAG-β-catenin K49R, had less interaction with RNAPII than wildtype β-catenin in response to H_2_O_2_ or PTEN KD. Also, FLAG-β-catenin K49R had reduced binding to the chromatin in response to PTEN KD. Our findings are unique from previous studies because we demonstrated that AKT-mediated phosphorylation of EZH2 at S21 (pS21-EZH2) drives EZH2 to interact with and methylate β-catenin, which enhances β-catenin’s and RNAPII’s binding to the chromatin. Based on our findings, we propose that the AKT-mediated phosphorylation of EZH2 results in EZH2’s regulation of β-catenin being dependent on the activity of EZH2, in contrast to other studies that have found that EZH2 regulates β-catenin independently of its catalytic activity (33). It should be noted that most of our models, like most colon cancers, have *APC* mutations resulting in constitutive activation of the WNT/β-catenin pathway and nuclear localization of β-catenin (27). We hypothesize that EZH2’s regulation of β-catenin in CRC is likely dependent on WNT pathway activation.

We also demonstrated that the AKT-EZH2-β-catenin axis regulates EZH2’s and β-catenin’s binding to chromatin and enrichment across the genome. While a phospho-mimetic EZH2 (EZH2 S21D) showed increased binding to chromatin, phospho-null EZH2 had reduced binding to chromatin relative to WT EZH2 following AKT activation. Our genome-wide binding profiles for HA-EZH2 and FLAG-β-catenin confirmed our hypothesis that EZH2 and β-catenin binding to the chromatin is dependent on EZH2 phosphorylation at S21 and EZH2 catalytic activity, respectively. Phospho-null EZH2 had a reduced number of called HA-EZH2 peaks in both EV and PTEN KD cells. Treating with an EZH2 inhibitor also drastically reduced the number of called FLAG-β-catenin peaks in both EV and PTEN KD cells. These findings are consistent with other findings in which EZH2 induced the binding of AR, cMyc, and P300 to the chromatin to mediate gene expression (34,35). While previous studies suggest that EZH2 methylation of other non-histone proteins leads exclusively to gene activation or repression (19,20), our findings propose a model in which EZH2-mediated β-catenin methylation increases β-catenin’s binding to the chromatin and fine-tunes β-catenin activity to both repress and activate gene expression. The exact mechanism of how methylation of β-catenin leads to gene activation and repression is unclear. It is possible that methylation of β-catenin represses gene expression by blocking β-catenin acetylation, which is associated with active gene expression, as has been reported previously (36). It is also possible that EZH2-mediated β-catenin methylation activates gene expression through increased recruitment of TCF1 and RNAPII to the chromatin. In breast cancer, phosphorylation of EZH2 by AKT was shown to attenuate EZH2’s catalytic activity by reducing its binding to the histones (37). While we see an increase in the binding of phosphorylated EZH2 to chromatin, we have not specifically examined how phosphorylation alters the binding to and/or methylation of histones.

Interestingly, our data suggest that AKT activation promotes β-catenin-mediated upregulation of EMT-related genes in an EZH2-dependent manner. We show that increased β-catenin enrichment over some EMT-related genes is associated with their increased gene expression in PTEN KD compared to EV cells. Treating cells with an EZH2 inhibitor resulted in reduced β-catenin enrichment across the genome. However, this finding was unexpectedly accompanied by further up-regulation of EMT-related genes. Related to this result, EZH2 inhibition increased expression of EMT-related genes in EV cells, which is similar to previous findings by other groups, and EZH2 is known to repress genes to maintain cell identity (29–31). For example, EZH2 preserves the identity of colon progenitor cells and maintains lineage differentiation commitment in the normal colon (38). During neuron differentiation, EZH2 also maintains cell identity (39). Additionally, a recent study showed that EZH2 maintained epithelial cell identity in non-small cell lung carcinoma cell, and inhibiting EZH2 activity induced cell transition to a mesenchymal phenotype (31). Therefore, it is possible that AKT-mediated EZH2 phosphorylation induces EZH2 to promote EMT in epithelial colon cells by interacting with and methylating β-catenin. However, inhibiting EZH2 activity also enhances colon epithelial cell plasticity by increasing the expression of EMT-related genes, which are usually repressed by PRC2. We speculate that pS21-EZH2 induces the expression of EMT genes through methylating β-catenin while inhibiting EZH2 attenuates the methylation of β-catenin and the histone-related repressive function of EZH2, which cumulatively results in increased EMT gene expression.

Based on our EZH2 CUT&RUN data, phosphorylation of EZH2 following AKT activation in PTEN KD cells greatly increased the number of genomic loci bound by EZH2. However, very few of these peaks were associated with changes in gene expression as they did not overlap with PTEN KD-regulated genes in our RNA-seq data. Most of the EZH2 peaks also did not overlap with FLAG-β-catenin peaks in PTEN KD cells in our ChIP-seq data. Because EZH2 inhibition blocked the PTEN KD-induced increase in β-catenin binding sites and altered expression of the PTEN-KD regulated genes, we conclude that EZH2-induced trimethylation of β-catenin modulates gene expression in PTEN KD cells. However, we suggest that most of the increased binding of EZH2 to chromatin following AKT-mediated phosphorylation on S21 is not regulated by its interaction with β-catenin and does not have a direct effect on gene expression. Future work will explore the mechanism and relevance of the increased binding of phosphorylated EZH2 to chromatin.

Overall, we purpose a model where activation of AKT induces EZH2 phosphorylation, which mediates β-catenin methylation. β-catenin methylation increases β-catenin’s interaction with TCF1 and RNAPII and increases β-catenin’s binding to the chromatin to regulate gene expression (Figure 8F). Extrapolation of the findings in this study suggests a need for the development of a context-specific strategy for the therapeutic targeting of EZH2 in CRC and that therapeutic inhibition of EZH2 in CRC should be considered with caution. The oncogenic role of EZH2 in cancers led to the development of many EZH2 inhibitors that are currently in clinical trials (40). However, our findings suggest that treatment with an EZH2 inhibitor will likely lead to different clinical outcomes depending on the mutations present in the specific CRC being treated. Our findings suggest that, in AKT-activated CRC, therapeutic approaches based on perturbation of the interaction between EZH2 and β-catenin or EZH2 and AKT will be more effective than targeting EZH2 activity as EZH2 inhibitors will block all activity dependent roles of EZH2, including EZH2’s methylation of histone and non-histone substrates.

## Materials and Methods

### Cell culture and treatments

All cell lines were maintained in a humidified atmosphere with 5% CO_2_. Our study included SW480, RKO, and HEK293T cells and human CRC organoids. SW480 cells were cultured in McCoy 5A media (Corning) and RKO cells were cultured in RPMI1640 media (Corning) supplemented with 10% FBS (Gibco). HEK293T cells were cultured in DMEM 1X with 10% FBS (Gibco) without antibiotics. CRC human organoids were cultured in organoid media (advanced DMEM/F12 (Gibco) supplemented with EGF (R&D Systems), Noggin (R&D systems), N2 supplement (17502048, Fisher), B27 supplement (17504044, Fisher), HEPES, and Penn/Strep) as in (41). All cell lines were purchased from the ATCC and authenticated and tested for Mycoplasma using the Universal mycoplasma detection kit (ATCC, 30-1012K) on January 30, 2023. All cells used in experiments were passaged fewer than 15 times with most being passaged fewer than 10 times. For hydrogen peroxide (H_2_O_2_) treatments, 30% H_2_O_2_ (Sigma) was diluted in PBS immediately prior to treatment at 250 mmol/L for 30 minutes at 37°C. Cells were starved in media lacking serum for 24 hours prior to treatment. GSK503 (Sigma, SML2718) and GSK690693 (Sigma, SML0428) were solubilized in DMSO (Sigma) prior to treatment. Treatment dosages and durations are defined in the figure legends.

### Generation of stable knockdown lines using viral shRNAs

For knockdown of EZH2 (Sigma, SHCLNG-NM, TRCN0000010474), PTEN (Sigma, SHCLNGNM_000314, TRCN0000002748), and empty vector (EV) TRC2 (Sigma, SHC201), the lentiviral shRNA knockdown protocol from The RNAi Consortium Broad Institute was used. Briefly, 4 × 10^5^ 293T cells were plated on day 1 in DMEM 1X containing 10% FBS. On day 2, cells were transfected with shRNA of interest, EV control, and packaging plasmids. On day 3, media was replaced with fresh DMEM containing 10% FBS. Approximately 24 hours later, media containing lentiviral particles was collected, and fresh DMEM + 10% FBS was added. The added media were collected 24 hours later and pooled with media harvested on day 4. The pooled media was then filtered using a 0.45 mm filter and concentrated using a Spin-X concentrator (Corning, #431490).

### Plasmids and transient transfections

HA-EZH2 plasmid mutant constructs were generated using site directed mutagenesis of pCMV-HA-EZH2 a gift from Kristian Helin (Addgene plasmid # 24230; http://n2t.net/addgene:24230; RRID:Addgene_24230 (42)). Wildtype and mutant constructs were then PCR amplified using primers to encompass HA-EZH2 + add NotI and PacI for subcloning into pQXCIH (Clontech). See Supplemental Table 1 for the primer sequences. FLAG-β-catenin plasmid constructs were purchased from Addgene (FLAG-β-catenin WT (Addgene plasmid #16828; http://n2t.net/addgene:16828; RRID:Addgene_16828), FLAG-β-catenin K49R (Addgene plasmid # 44750; http://n2t.net/addgene:44750; RRID:Addgene_44750), FLAG-β-catenin K19R (Plasmid #Addgene plasmid # 44749; http://n2t.net/addgene:44749; RRID:Addgene_44749), FLAG- β-catenin K19R/K49R(Addgene plasmid #44751; http://n2t.net/addgene:44751; RRID:Addgene_44751)). All FLAG- β-catenin constructs were a gift from Eric Fearon (26,43). Transient transfection was performed with Lipofectamine 3000 (Invitrogen) per the manufacturer’s protocol.

### Antibodies

For Western blot of endogenous proteins, anti-EZH2 [Cell Signaling Technology (CST, #5246, 1:1,000], anti–β-catenin (CST, #8480, 1:1,000), anti–TCF1 (CST, #2203, 1:1,000), anti-Rpb1 CTD (4H8) (CST, #2629, 1:1,000), anti-Vimentin (CST, #5741, 1:1,000), anti-Phospho-Akt (Ser473) (CST, #4060, 1:1,000), anti-total AKT (CST, #4691,1:1,000), anti-TCF1 (SC, sc-271453, 1:1,000), anti-H3K27me3 (CST,#9733, 1:1000), anti-H3(CST, #4499, 1:1000) anti-GAPDH (CST, #5174,1:1,000), anti-Phospho-Akt Substrate (RXXS*/T*) (CST, #9614, 1:1000), and anti-FLAG (sigma, F3165) antibodies were used. For IPs, EZH2 (CST), TCF1 (CST), β-catenin (CST), and FLAG (sigma) antibodies were used.

### Nuclear Immunoprecipitations (IPs)

3.5 × 10^6^ or 1.2 × 10^6^ cells were cultured in 150 mm plates for approximately 72 hours or 96 hours, respectively. Cell pellets were used to perform nuclear extraction using CEBN [10 mmol/L HEPES, pH 7.8, 10 mmol/L KCl, 1.5 mmol/L MgCl_2_, 0.34 mol/L sucrose, 10% glycerol, 0.2% NP-40, 1X protease inhibitor cocktail (Sigma, P5726), 1X phosphatase inhibitor (Thermo, 88266), and N-ethylmaleimide (Acros organics, 128-53-0)] and then washed with CEB buffer (CEBN buffer without NP-40) containing all the inhibitors. To extract the soluble nuclear fraction, after washing the cell pellets with CEBN buffer, they were resuspended in modified RIPA (50 mM Tris pH7.5, 150 mM NaCl, 5 mM EDTA, 50 mM NaF, all inhibitors), sonicated using Bioruptor® Pico (Diagenode) and rotated at 4°C for 60 minutes. The nuclear extract was rotated with an antibody and protein A/G magnetic beads overnight. The next day, beads were washed, and proteins were eluted and analyzed by Western blot. For some IPs, RNAseA (25 μg/ml) was added during the overnight incubation.

### Chromatin extraction

3.5 × 10^6^ or 1.2 × 10^6^ cells were cultured in 150 mm plates for approximately 72 hours or 96 hours, respectively. Cell pellets were used to perform nuclear extraction using CEBN and then washed with CEB buffer containing all the inhibitors. To extract the soluble nuclear fraction, after washing the cell pellets with CEB buffer, they were resuspended in soluble nuclear buffer (2 mmol/L EDTA, 2 mmol/L EGTA, all inhibitors) and rotated at 4°C for 30 minutes. The remaining cell pellet, i.e., the total chromatin fraction, was lysed using 4% SDS and a qiashredder (Qiagen) and analyzed by Western blot.

### Chromatin immunoprecipitation sequencing

Chromatin immunoprecipitation (ChIP) was performed using anti-FLAG antibody (Sigma) and the Diagenode iDeal ChIP-seq Kit (C01010055, for transcription factors) as per the manufacturer’s protocol. Libraries were generated for sequencing using the NEBNext Ultra II DNA Library Prep kit for Illumina (NEB) as per the manufacturer’s protocol, followed by sequencing.

### CUT&RUN

CUT&RUN was performed with anti-HA antibody (HA Tag CUTANA, 13-2010) and a commercial kit (EpiCypher CUTANA pAG-MNase for ChIC/CUT&RUN, 15-1116) as per the manufacturer’s protocol. 5 × 10^5^ cells were used for CUT&RUN, and 5 ng of the purified CUT&RUN DNA was used for construction of multiplexed libraries with a NEBNext Ultra II DNA Library Prep kit, followed by sequencing.

### RNA sequencing

RNA extraction was performed using the RNAeasy mini kit (Qiagen, 74104). Libraries were generated for sequencing using the NEBNext Ultra II DNA Library Prep kit for Illumina (NEB) as per the manufacturer’s protocol, followed by sequencing.

### Migration assays

5 × 10^4^ SW480 cells in serum-free media were plated into transwell in 24-well plates (Corning #40578) for 48 hours with media containing 10% FBS at the bottom. Transwell inserts were stained using Hema 3 Stat Pack (Thermo Fisher Scientific #123-869). Migration inserts were randomized prior to manual quantification and the outer 5% of the inserts were not included during quantification to reduce edge-effect bias. All images were taken on an EVOS FL Auto microscope.

## Acknowledgements

We would like to thank the Indiana University Center for Genomics and Bioinformatics for their assistance with library preparation and sequencing.

## Funding

This work was funded, in part, with a Core Pilot Grant [to H.M. O’Hagan] from the Indiana Clinical and Translational Sciences Institute funded, in part by Grant Number ULITR002529 from the National Institutes of Health (NIH), National Center for Advancing Translational Science, Clinical and Translational Sciences Award. This work was also supported by a Research Enhancement Grant and Elwert Award [to H. M. O’Hagan] from the IUSM and pilot funding [to H. M. O’Hagan] from the IUSCCC P30 Support Grant (P30CA082709). The content is solely the responsibility of the authors and does not necessarily represent the official views of the NIH or IUSM. Additional research funding was provided by Van Andel Institute through the Van Andel Institute - Stand Up to Cancer Epigenetics Dream Team [to H. M. O’Hagan]. Stand Up to Cancer is a division of the Entertainment Industry Foundation, administered by AACR. A. Ghobashi was supported by the Doane and Eunice Dahl Wright Fellowship generously provided by Ms. Imogen Dahl.

## Notes

### Competing Interest Statement

The authors have declared no competing interest.

### Summary of Updates

The first author's name was updated to include his middle initial.

