## Supplemental methods and Supplemental Table 1 for "Activation of AKT induces EZH2-mediated β-catenin trimethylation in colorectal cancer"

Base editing of the *PIK3CA*

To generate sgRNA expressing plasmids, E453K or E453K specific oligos were annealed and cloned into pLenti-U6-tdTomato-P2A-BlasR (LRT2B), a gift from Lukas Dow (Addgene plasmid #110854; http://n2t.net/addgene:110854; RRID:Addgene_110854 (1)). SW480 cells were then transfected with E453K, E453K, or empty vector LRT2B plasmids and pLenti-FNLS-P2A-Puro, a gift from Lukas Dow (Addgene plasmid #110841; http://n2t.net/addgene:110841; RRID:Addgene_110841) at a 1:3 ratio of sgRNA:pLenti-FNLS using lipofectamine 3000. The next day media was changed to fresh media containing puromycin. Following puromycin selection, cells were grown in MEK inhibitor (25 nM Trametinib) for 7 days to select for cells containing mutant *PIK3CA*. To check for successful base editing, genomic DNA was isolated using the DNeasy kit (Qiagen), PCR amplified using E545K or E453K AP PCR primers, and purified PCR products (Qiagen PCR purification kit) were sequenced by Sanger sequencing using the respective sequencing primers. See Supplemental Table 1 for the oligo and primer sequences.

Sequencing Analysis

Sequencing read quality control for all samples was assessed with FastQC (v0.11.5). For RNA-seq analysis, read alignment was performed using STAR (2) (v2.6.1a) against the hg38 reference genome, raw read counts for genes were obtained with STAR (v1.24.2), and Deseq2 (3)(v3.24.0) was used for differential expression analysis. Gene Set Enrichment Analysis using the Molecular Signatures Database collections and database were run with clusterProfiler (4)(v3.12.0) and volcanoplot was visualized with EnhancedVolcano. For ChIP-seq and CUT&RUN analysis, read alignment was performed using Bowtie2 (5)(v2.3.2) against the hg38 reference genome, peaks were called with MACS (6)(v2.1.0), and intersecting peak regions between replicates and different samples were uncovered with Bedtools (7)(v2.26.0). Heatmaps and Counts Per Million normalized bigWigs were created using deepTools (8)(v3.1.3) bamCoverage, computeMatrix, and plotHeatmap. Gviz (9)(1.28.0) was utilized for creating gene tracks. The correlation between EZH2 and PTEN expression in CRC was performed using UCSC xenabrowser (10)

| Primers/oligos | Sequences |
|---|---|
| E545KP3 | P-5'-CACCGCGTCAGTGATTTCAGAGAG 3' |
| E545KP4 | P-5'-AAACCTCTCTGAAATCACTGACGC 3' |
| E453KP3 | P-5'-CACCGTCTTCTAATCCATGAGGTAC 3' |
| E453KP4 | P-5'-AAACGTACCTCATGGATTAGAAGAC 3' |
| E545K_seq F1 | CATCTGTGAATCCAGAGGGGAA |
| E545K_seq R1 | TGCTGAGATCAGCCAAATTCA |
| E453K_seq F1 | GGGGAAAAAGGAAAGAATGGGC |
| E453K_seq R1 | GAGAGAAGGTTTGACTGCCA |
| E545K AP_F1 | AGCTTTGCAGGGATCATAAGG |
| E545K AP_R1 | CGTATCACCAACAGCAGGGTA |
| E453K AP_F1 | TACCTTGGGAGAGCTTCAGGA |
| E453K AP_R1 | ACTCAGTGATTTGCCTTACCAGT |
| 5' EZH2 with 1X HA tag | CTATGCATACCCATACGATGTTCCAGATTACGCTATGGGCCAGACTGG GAAGAAATC |
| 3' EZH2 adds Pac1 site | GTGACATTAATTAATTATCAAGGGATTTCCATTTCTCTTTCG |
| 5' PCR sewing primer for EZH2 point mutation in S21A | CGGAAGCGTGTAAAAGCAGAGTACATGCGACTG |
| 3' PCR sewing primer for EZH2 point mutation in S21A | CAGTCGCATGTACTCTGCTTTTACACGCTTCCG |
| 5' PCR sewing primer for EZH2 point mutation in S21D | CGGAAGCGTGTAAAAGACGAGTACATGCGACTG |
| 3' PCR sewing primer for EZH2 point mutation in S21D | CAGTCGCATGTACTCGTCTTTTACACGCTTCCG |

**Supplemental Table 1. Sequences of primers and oligos used for PIK3CA base**
